# Adapting a gas chromatograph for use as a testing chamber for thermal ecology experiments

**DOI:** 10.1101/2025.05.22.655604

**Authors:** Gabriel Zilnik, Miles T. Casey, Paul V. Merten, Scott Machtley, James R. Hagler

## Abstract

1. Temperature impacts many aspects of species biology and ecology. A continuing struggle for studies in thermal ecology is accurate assessment of critical thermal maxima *CT*_max_. Identifying when loss of equilibrium (LOE) occurs has been criticized for being too subjective. This is particularly true of small organisms, such as insects, where the loss of coordination can be difficult to observe. As such, ecologists have often used lack of movement as a proxy for LOE.
2. Here, we designed, tested, and present a guide to recycling surplus gas chromatographs for use as a thermal chamber that allows accurate and high throughput assessment of *CT*_max_ at relatively low cost.
3. We found the GC to be an adequate heating chamber for thermal experiments. Installation of a rotating rack that can hold glass observation vials allows for rapid identification of loss of equilibrium in subjects. We evaluated the *CT*_max_ of a common generalist predator in the Arizona cotton agroecosystem, *Collops vittatus*. Additional tests of static heat exposure also revealed that this chamber can be used for assessing the impacts of heat stress on predatory behavior.
4. We hope to encourage other ecologists to use this guide to recycle surplus laboratory equipment for use in thermal ecology studies. We believe that our thermal insect carousel can be used for *CT*_max_, lethal temperature, and behavioral bioassays.

## INTRODUCTION

Thermal performance influences organismal physiology, behavior and fitness (Angilleta 2009; Huey and Kingsolver 2011). Understanding the thermal environment of species is crucial to understanding their distribution and abundance (Jiang and Morin, 2004; Angiletta et al, 2006; Niehaus et al, 2012; Faillace et al, 2021). One key measure of an organisms response to it’s thermal environment is the critical thermal maximum (*CT*_max_), the point at which it loses locomotory control and its ability to escape from conditions that will lead to death (Cowles and Bogert 1944). This measure has been used for nearly a century to study the effects of temperature on organisms (Lutterschmidt and Hutchison, 1997; Rohr et al, 2018). To investigate this relationship rigorously, research requires precise control over thermal conditions, which is often difficult to accomplish in natural settings due to environmental variability. Laboratory-based testing offers customizable experimental environments that are repeatable and stable.

Despite their utility many existing thermal chambers are limited in scalability, flexibility, or cost-effectiveness. Commercial thermal chambers range from US$5,000 to >US$20,000, placing them out of reach for researchers with limited funding. Low-cost solutions, such as water baths, can often struggle with spatial temperature heterogeneity if not actively mixed (Huey and Stevenson 1979; Lutterschmidt and Hutchison 1997). A potential solution has been modification of consumer equipment to achieve lower cost experimental environments. Greenspan et al. (2016) were able to achieve accurate and repeatable ramping and cooling to within ±0.5 °C with inexpesive modifications to a commercially available constant temperature chamber. Here, we describe the conversion of a surplus Gas Chromatograph (GC) into a thermal insect carousel (TIC), that has greater than five times the thermal accuracy of their chamber. Our prototype, constructed at minimal cost (<$100US beyond the free base unit) from a retired GC which only had a functional oven demonstrates how obsolete laboratory equipment can be transformed into precision research tools. Researchers with limited research funds could repurpose a surplus GC and adapt it for thermal ecology experiments for far less capital than new specialized equipment, while also extending the functional lifespan of existing laboratory equipment.

We validated the TIC’s utility through two proof-of-concept experiments with the predatory beetle *Collops vittatus* (Coleoptera: Melyridae). The first was a dynamic assessment of the *CT*_max_. The second was a static experiment to determine the impact of heat stress on the feeding behavior of *C. vittatus.* We demonstrate that the TIC combines the precision of commercial systems with the affordability of DIY approaches, while introducing unique capabilities, like high-throughput sampling.

## MATERIALS AND METHODS

### TIC Construction

Measurements are given in metric when metric components are used, but otherwise Society of Automotive Engineers (SAE) measurements are provided due to hardware that was available on hand or could be obtained from a local hardware store. Metric hardware could be swapped as we are providing a general plan for converting a GC to a thermal insect carousel (TIC). Where SAE are used, we have provided the nearest metric equivalent in brackets (e.g. 1 inch [25 mm], 0.375 inch [9 mm or 10 mm]).

1. *Removing GC components*

a. The TIC was created from a malfunctioning HP Agilent 6890 series gas chromatograph (GC) (Agilent Technologies, Santa Clara, CA).
b. The column equipment, column heaters, tubing, gauges, solenoid switches and door were removed.
c. As each component was removed, we activated the GC and tested the oven to ensure the component removal did not alter its functionality.

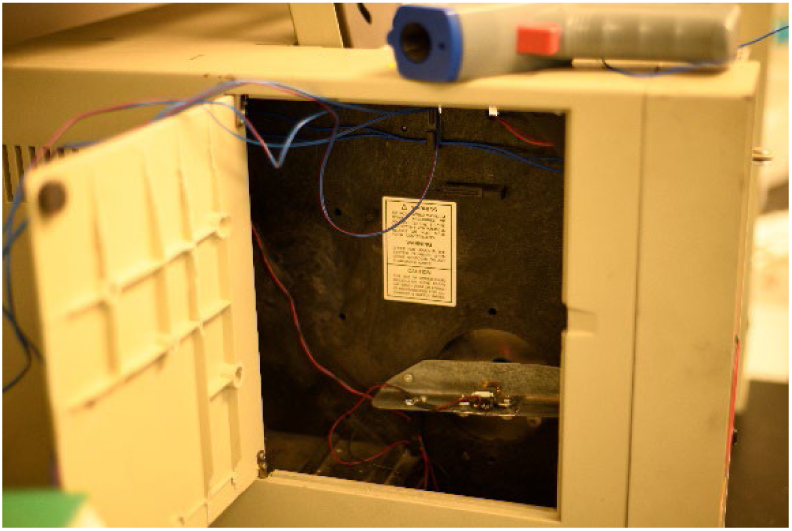
d. We replaced the original control face plate with 20 gauge [0.813 mm] sheet metal, and use its surface to mount thermometers, and the controls for lighting and an electric rotary motor (detailed below in Steps 3 & 4).
e. A new door was fabricated using 0.125 in [3 mm] steel plate. A square opening was cut out and a glass pane fitted over it with silicone adhesive. This change permitted easy viewing of subjects.

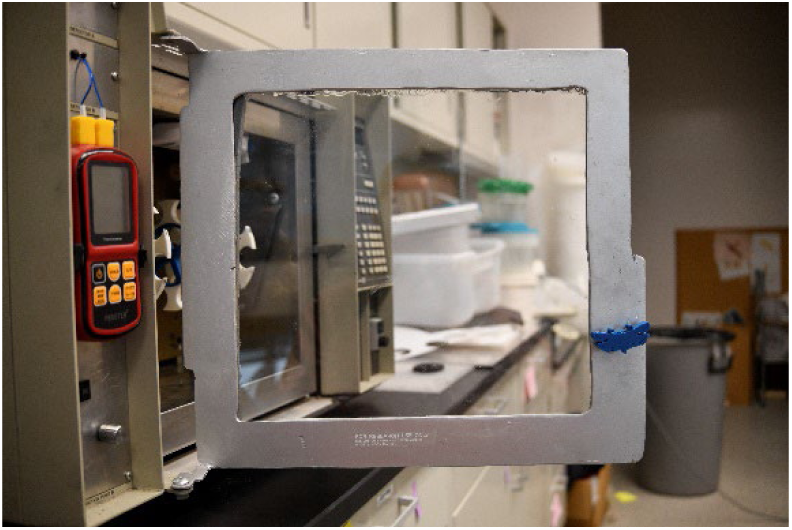
f. Heat loss was prevented by sealing the door frame with 0.75 in [19 mm] high-density foam weatherstripping (Product: 02311, M-D Building Products, INC, Oklahoma City, OK, USA).

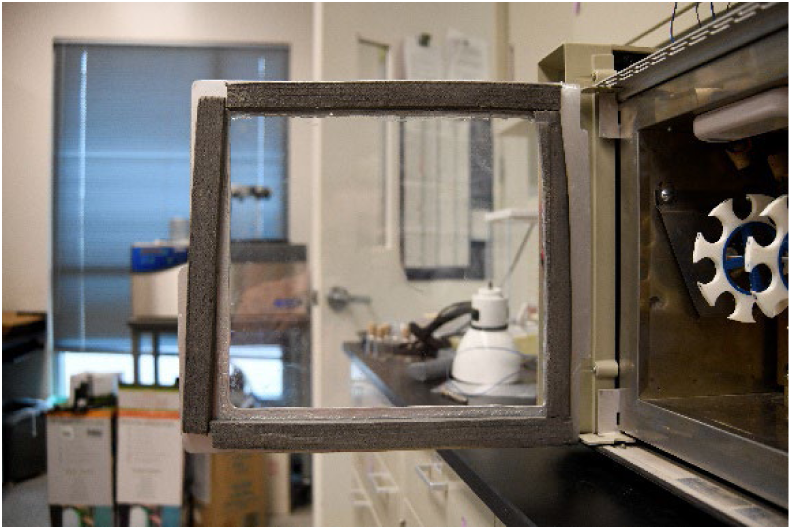
2. *Mounting motor and controls*.

a. To create mount for an electric rotary motor (1203BB, UXCELL, Hong Kong), a section of sheet metal was attached to the side of the oven chamber. Aligned opening were drilled into both surfaces. These had an aperture of 0.3125 in [8 mm] and were fitted with a 6 mm nylon snap bushing into which a 6×13×5 mm ball bearing (Part: F686ZZ, Uxcell, Hong Kong) was inserted.
b. The micro gear motor was then mounted to the sheet metal using and angle iron that could act as a support for the motor.

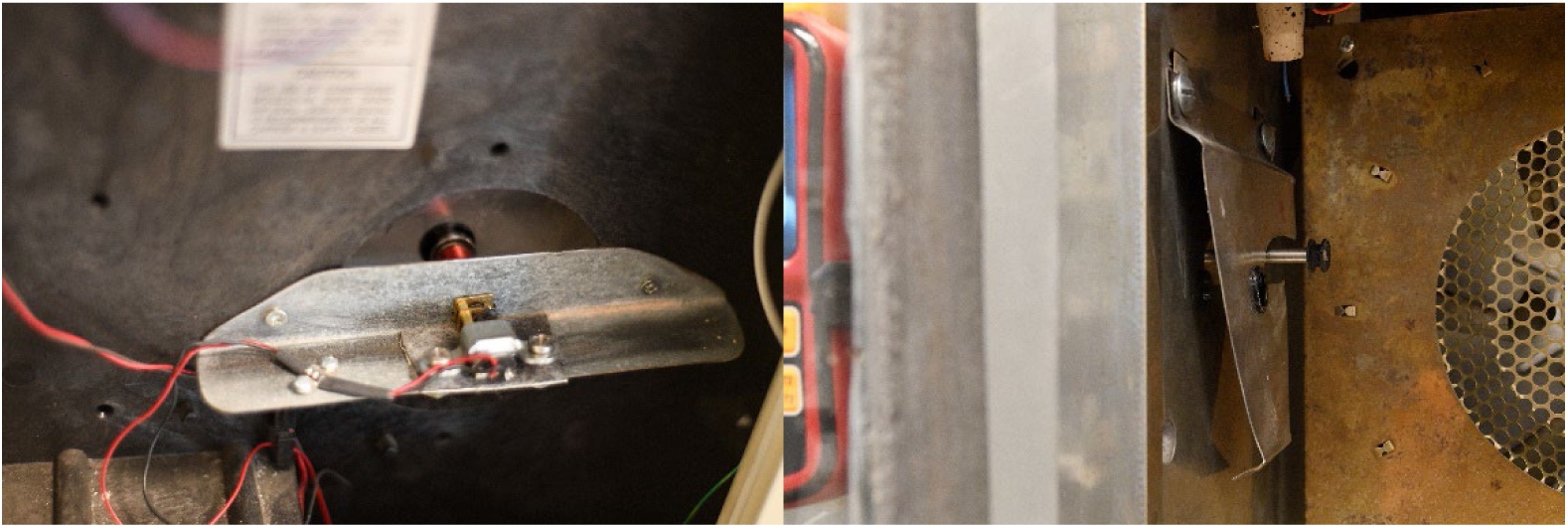
c. The motor was attached to a 6 mm diameter drive shaft.
d. We secured a plastic screen door roller to the drive shaft to act as a pulley.

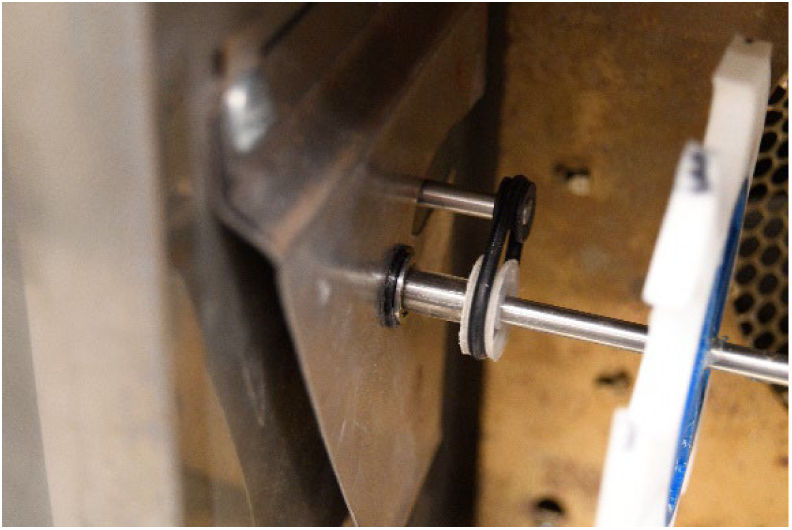
e. After locating a 5-volt power source on the motherboard, we wired it to a motor speed controller (Model: 1203BB 6V 12V 3A 80W DC Motor Speed Controller, Ledomo, China) mounted to the front of the GC.

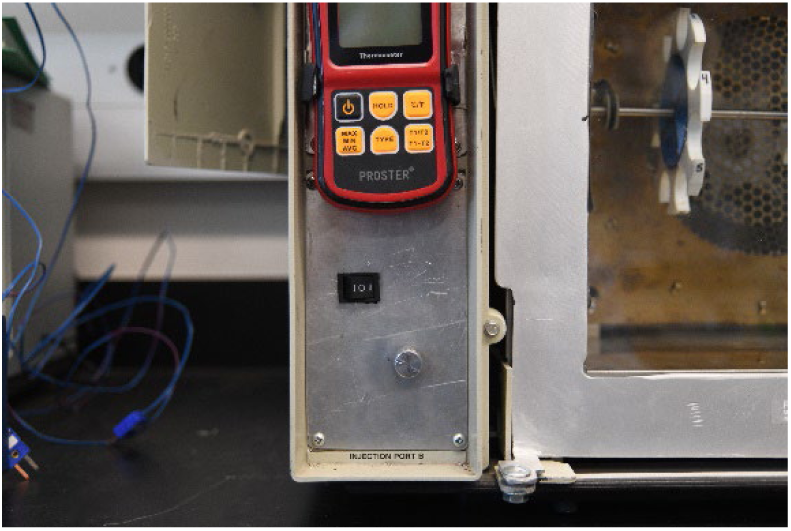
3. *Mounting Thermometer and Thermocouples*

a. Sheet metal was used to create a bracket to hold a battery powered digital thermometer (Model: 4333090752, Proster Trading Limited, Hong Kong). The thermometer was later replaced with a more accurate (±0.01 °C vs ±0.5 °C) and T-type thermocouple system (Model: TC-2000, Sable Systems International, North Las Vegas, Nevada, USA). The TC-2000 is too large to mount to the TIC so we place it next to the TIC on the bench.

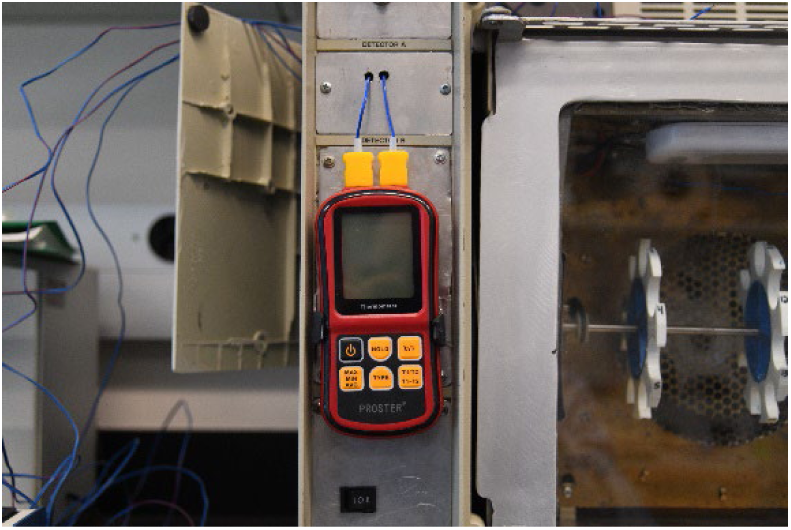

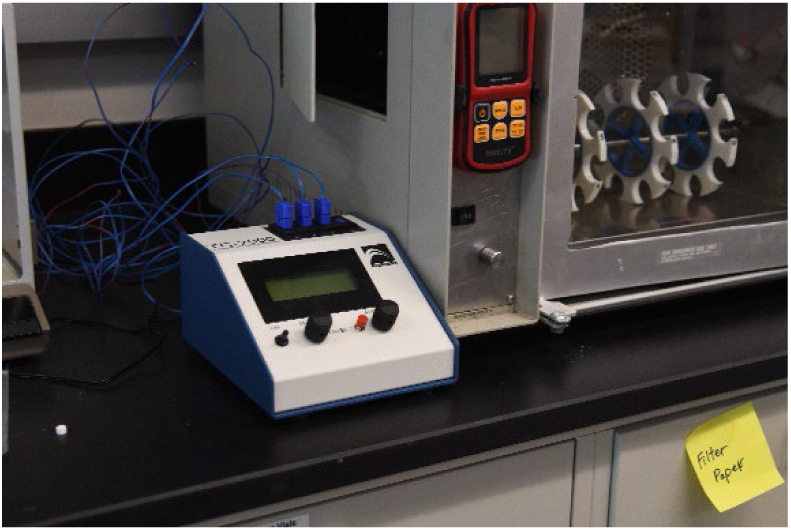
b. The top plate of the GC was fitted with rubber grommets in place of the removed column heaters, allowing us to securely pass the thermocouples into the oven.
c. We mounted two scintillation vials (5 mL and 20 mL) on the top surface inside the oven and inserted two K-Type thermocouples (later T-Type) through cork vial stoppers to monitor the temperature inside of the vials, which served as a representation of the temperature inside of vials on the rotary racks.

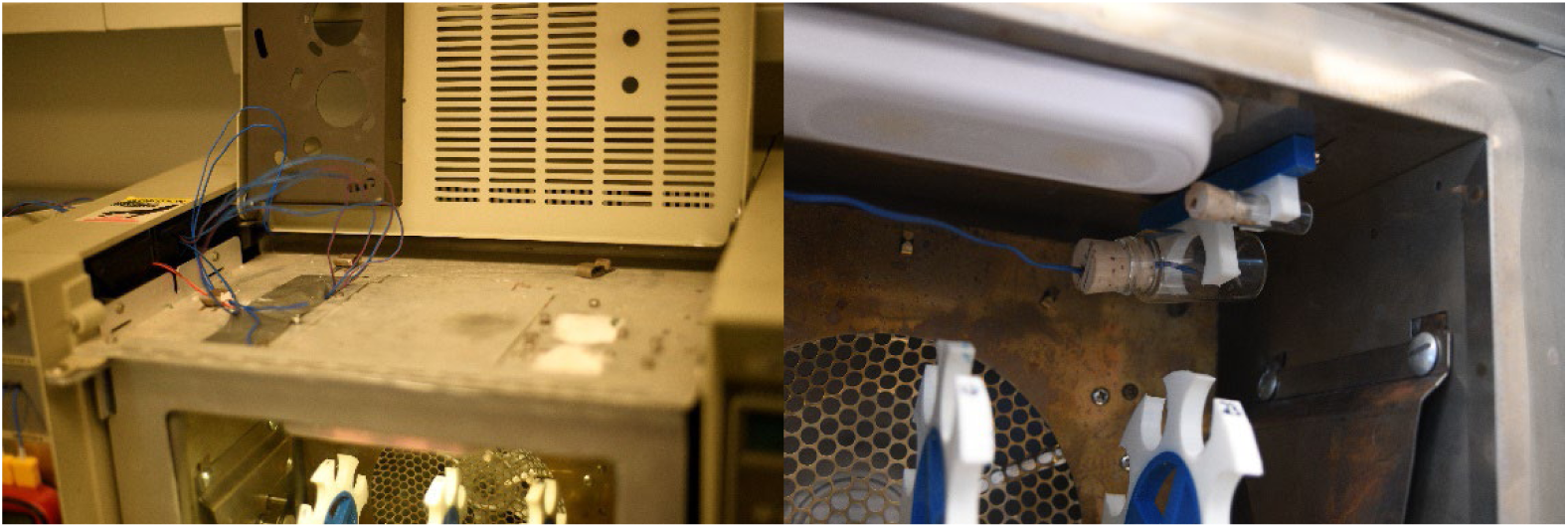
4. *Lighting*

a. We mounted a toggle switch to the top section of the front face plate and ran power from the speed controller to the toggle switch.
b. Power was routed along the same path as the thermocouples to a 6-volt, battery powered LED light (Model: 4141, Walmart Great Value, Arkansas, USA).
c. To bypass the battery, we soldered the electrical wires to terminals inside of the light.
5. *Attaching and Mounting Rotary Racks with Vial Holders*

a. Rotary rack wheels were designed using free online software (www.tinkercad.com; Supplementary Files 1-2).
b. Wheels were constructed with a 3D printer (UltiMaker S5, Ultimaker B.V., Geldermalsen, Netherlands) using central spokes made of PLA (Ultimaker B.V.) and outer wheel made of TPU 95A (NinjaFlex Snow White, NinjaTek, Lititz, PA).

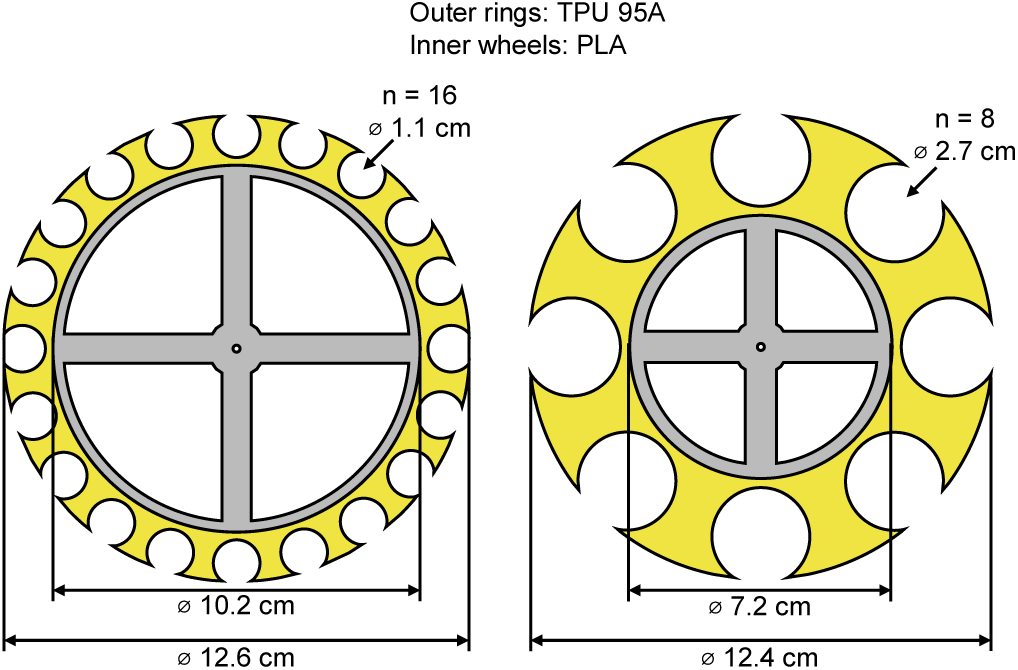
c. Wheels were friction fit to the drive shaft and arranged as either three wheels holding eight 20 mL vials each (total = 24 vials) or five wheels holding 20 5 mL vials each (total = 100 vials).
d. On the drive shaft, from left to right, we installed: 6 mm length of vinyl tubing (serving as a bearing retainer), 6 mm bearing, 13 mm length of vinyl tubing, screen door roller, 3 scintillation vial wheels (spaced evenly apart), 13 mm length of vinyl tubing, 6mm bearing, and 6mm length of vinyl tubing.

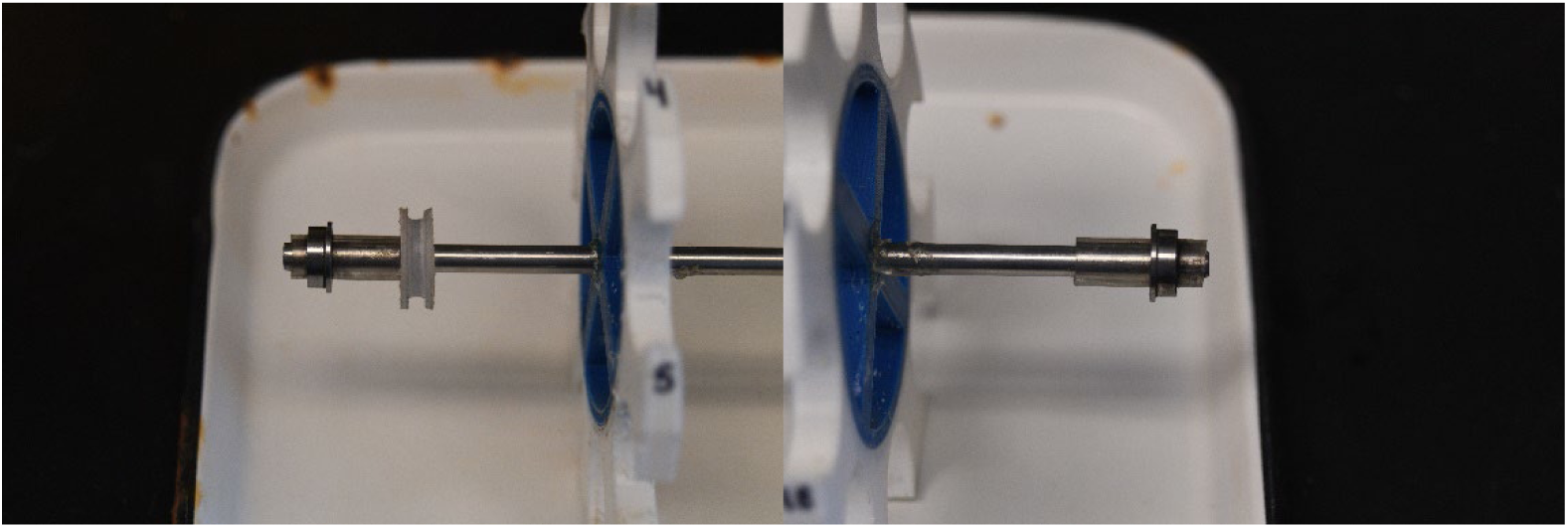
e. A 20 gauge [0.813 mm] thick sheet metal was cut into a trapezoidal shape and bent to hold the plate approximately 13 mm away from the wall. The stand-off from the side of the oven allows for removal and replacement of the entire rotary rack assembly by compressing one plate at a time. This capacity for easy removal allows specimen bottles to be loaded and unloaded outside of the TIC.
f. The top of the sheet metal was bolted to the sides of the oven with 0.25 in [6mm] bolts.
g. Holes were drilled into each plate that aligned with the center of the oven, and these were fitted with 6mm bearings and snap bushings.

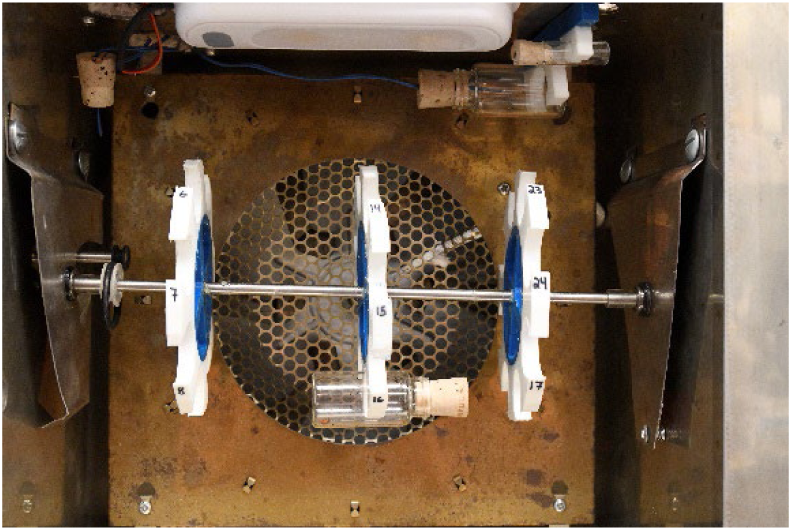
h. We drilled an additional hole on the left plate to allow for the motor axel to come into the oven.

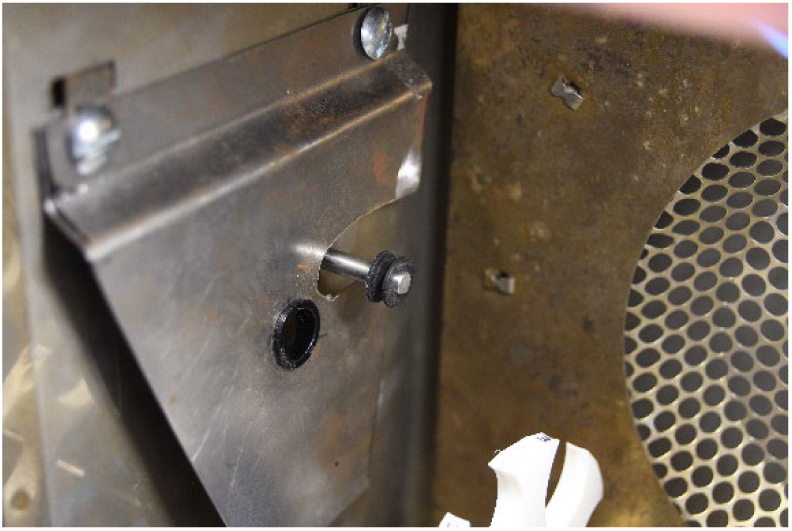
i. Using a 1 inch [25 mm] rubber o-ring (35771B, Danco Incorporated, Irving, TX, USA) to attach the pair of screen door rollers, we connected the motor drive shaft to the rotary rack shaft.

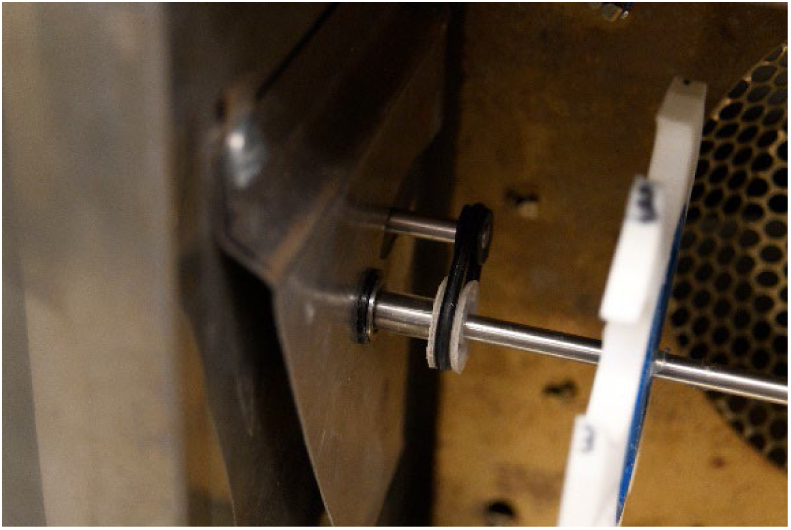

### Operation

Once the TIC is turned on, via the GC master power switch, it will begin the standard initial diagnostic procedures. (During this period, test animals can be loaded into the rotary rack.) The rack shaft can then be attached to the motor and rotations started. The light inside of the chamber can also be switched on. While the insects acclimate to this new environment, the TIC temperature can be programmed via the GC control panel. For dynamic experiments, we program an acclimation period of 30 min at 30 °C, then have a ramp temperature of 0.5 °C · min^-1^, with a maximum temperature of 60 °C to be held indefinitely. This final setting was chosen as an end point because no insects can survive protracted exposure to this temperature. For static experiments users can either choose to ramp to the static temperature or not. In between runs on the TIC, we turn off the oven function and we leave the door open. The small size of the chamber allows for rapid equalization of the oven chamber’s temperature to ambient ±0.5 °C.

### Thermal Stability Test

We programmed the TIC to hold a single temperature for 8 h. We evaluated stability between 30 °C and 60 °C in 5 °C increments. We recorded the temperature using the TC-2000 and four T-type thermocouples from four locations inside of the oven: 1) The top mounted vial detailed above; 2) A vial on the center rotary rack wheel held in the position closest to the door; 3) A vial on the right wheel held in the position furthest from the door; and 4) A vial on the left wheel held in the position closest to the bottom of the oven. Temperatures were recorded every hour.

### Insect Bioassays

To demonstrate the functionality of the TIC for studies of thermal ecology, we conducted two standard experiments for this field using the predatory beetle, *Collops vittatus*. These were collected with sweep nets from alfalfa fields at the Maricopa Agricultural Center research farm (Maricopa, AZ, USA) on 29 April 2025. The recorded temperatures for that day were T_max_ = 30.3 °C; T_min_ = 10.9 °C; and T_avg_ = 21.3 °C. The preceding 14 days had an average (±SD) T_max_ = 30 (±3.9) °C; T_min_ = 12 (±3.1) °C; and T_avg_ = 21.6 (±2.7) °C. Beetles were separated by sex into assorted 11.5 cm x 7.5 cm (diameter x height) plastic deli containers (Model: S-22770, ULINE, Pleasant Prairie, WI, USA). The beetles were provided a cotton ball moistened with water. We either subjected the beetles to CT_max_ assay immediately after collection and separation, or starved them for 24 h and subjected them to the heat stress feeding behavior assay.

The first bioassay was a determination of the beetles’ *CT*_max_. Twelve males and twelve females were placed individually into 20 mL scintillation vials, closed with a cork stopper, and loaded into the TIC. The motor was engaged at a constant speed of 2 RPM. The temperature protocol was as follows: 30 min at 30 °C, then 0.5 °C · min^-1^ until we observed loss of equilibrium (LOE). We determined LOE to be when the insects could no longer maintain their hold on the vial wall and no longer right themselves after falling from the walls. The *CT*_max_ was calculated as the arithmetic mean of LOE for males, females, and both combined. We evaluated differences between sexes via ANOVA.

The second bioassay conducted was an examination of the impact of heat stress on the beetles’ feeding behavior, we took twenty individuals of each sex and separated them into two treatments: 1) no-stress which were held at ambient temperature (27 ±2 °C); and 2) heat-stress which were held at 46 °C inside of the TIC. We chose 46 °C as that is, generally, a peak summer temperature for midday during the height of the cotton growing season in AZ when potential prey items would be present (Luttrell et al, 2015; Allen et al, 2018). Each treatment group was held at the acute exposure temperature for 4 h then allowed to recover at ambient temperature (24 ±1 °C) for 1 hour inside of an individual 100 mm diameter petri dish. After 1 h, individuals were presented with a sheet containing between 44 and 76 *Lygus hesperus* eggs. We recorded the proportion of eggs consumed during 1 h. We evaluated the difference in egg consumption between sex, treatments and their combinations via Wilcoxon rank-sum test.

### Statistical Evaluation and Visualization

All statistical tests were done in R version 4.4.1 (CRAN 2024). Data were visualized in *ggplot2* (Wickam 2016).

## RESULTS

### Thermal Stability

Over the course of 8 hs the static temperature test showed the TIC stayed within −0.3 to +1.1 °C of the set temperature (Fig 1A). We found the average fluctuation between readings was 0.06 °C (SEM: 0.02 °C, Fig 1B).

**Figure 1.**
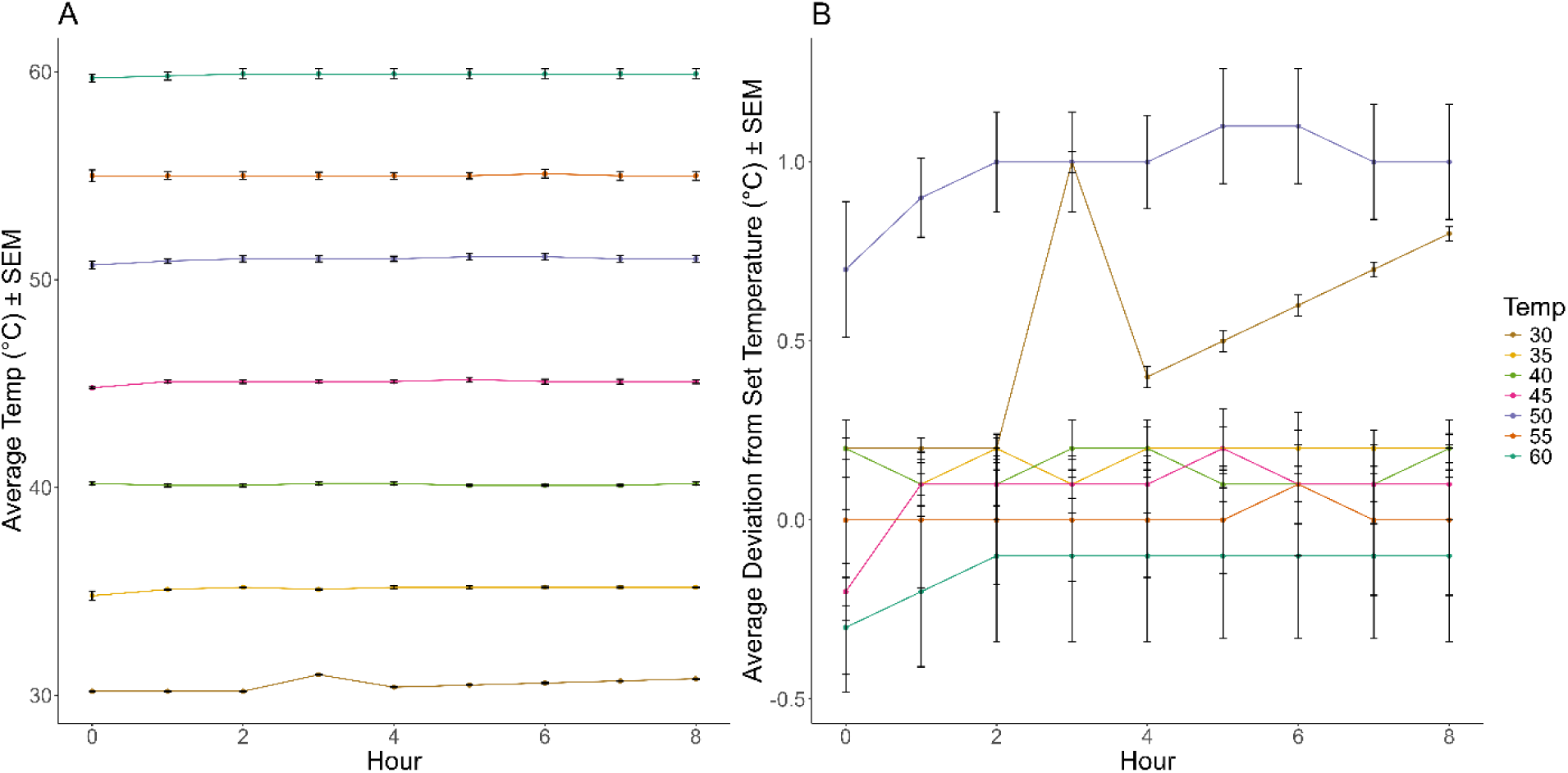
Average temperature (°C) over eight hours at set points between 30 °C and 60 °C (A). Deviation from the set point at each hour (B).

### Insect Bioassays

The *CT*_max_ for *C. vittatus* was 52.1 °C (SEM: ±0.35 °C). Males had a *CT*_max_ of 52.0 °C (SEM: ± 0.24 °C, n = 11) and females were 52.2 °C (SEM: ± 0.23 °C, n = 12). There was no difference between male and female *CT*_max_ (F_1, 21_ = 0.314, p = 0.581). Heat stress was found to impact feeding behavior. The stress was insufficient to induce mortality in any of the treatment groups but did reduce the number of prey eaten (Wilcoxon rank-sum W = 375.5, p < 0.001, Fig. 2A). Both males and females consumed fewer prey items when heat stressed (W_male_ = 87.5, p = 0.005; and W_female_ = 102, p = 0.003; Fig 2B). We observed no differences between males and females within treatments (No Stress: W = 46.5, p = 0.390, Heat Stress: W = 29.5, p = 0.062; Fig. 2B).

**Figure 2.**
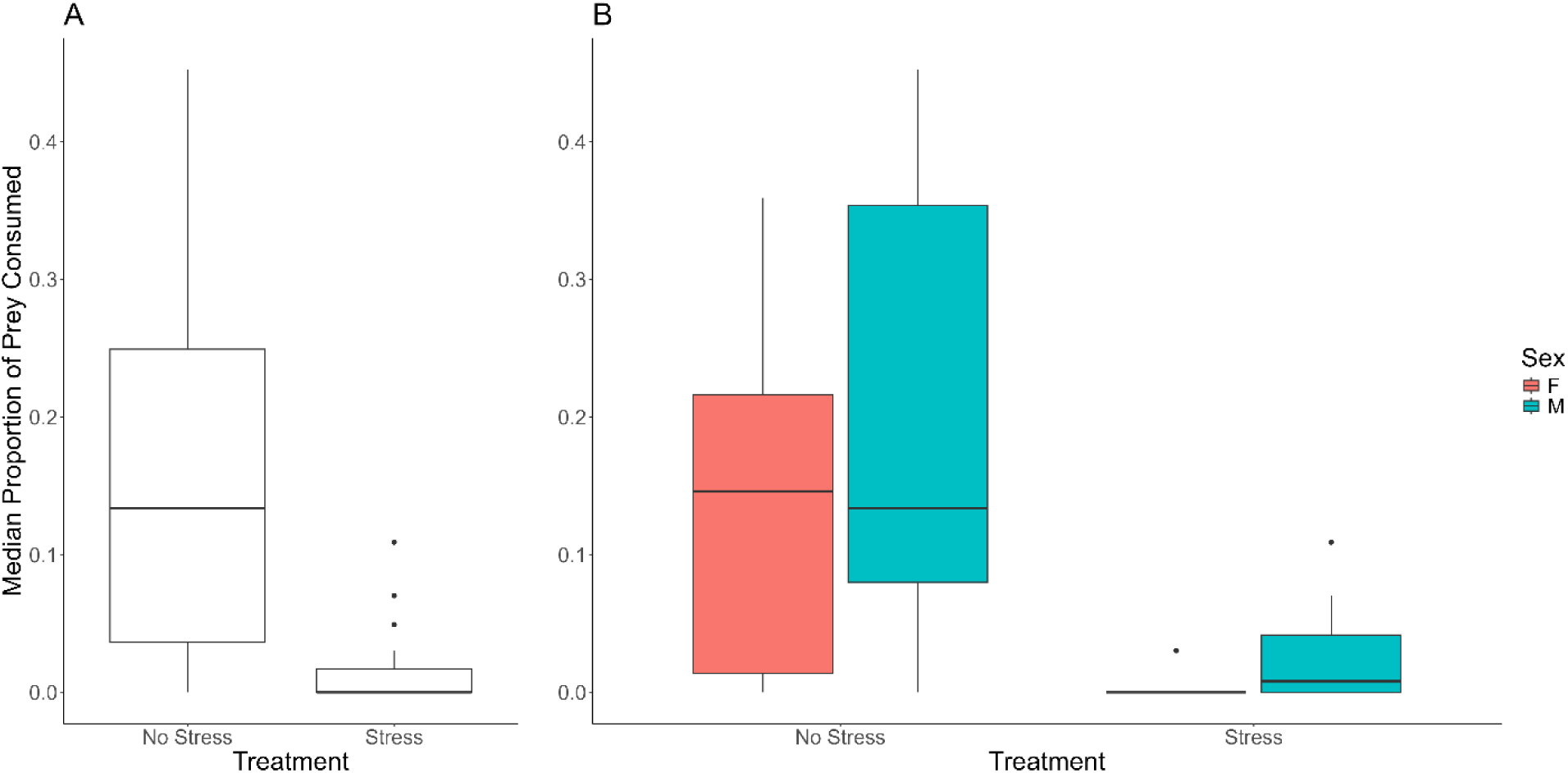
Median proportion of prey consumed across treatments (A) and between males and females within treatments (B). Fewer prey were consumed after the heat stress treatment, but no differences were observed between males and females within treatments.

## DISCUSSION

We repurposed a surplus GC into a versatile thermal testing chamber for ecolological experiments. This conversion takes advantage of the GC’s precision oven and programmability, which maintains stable temperatures (± 0.06 °C). A preliminary search on secondary markets (e.g. eBay) found that Agilent 6890 series GCs can be found for as little as $800US. Costs could be further reduced as universities and other laboratory facilities may have disused or non-functional GCs in their surplus equipment storage facilities. If the oven controls are operating, the absence of the analytical tools does not impact the functionality of the TIC. This approach can transform obsolete or malfunctioning equipment into a precision thermal chamber at a fraction of the cost of commercial thermal chambers. In our case, a malfunctioning GC was acquired at no cost and required <$100US in modifications to achieve performance comparable to commercial chambers. Furthermore, this approach aligns with growing efforts to reduce scientific waste through equipment reuse (Paes et al, 2017; Monson et al, 2025)

Another benefit of our design is that it is modular. The wheels of the rotary rack can be fabricated to hold various size vials depending on insect size and vial availability. For our use, we 3D-printed the wheels for the two most common vial sizes we already had in our facility. Using free online computer-aided design software, wheels can be designed to hold whatever size vials a researcher has on hand that would fit within the oven chamber. While we include plans for a wheel set capable of holding 100 vials of smaller insects, we found that recording data for 24 specimens approached the upper limit for a single observer. If higher throughput is desired, then multiple observers would be recommended to ensure that test subjects reaching their *CT*_max_ are not overlooked. When additional funds became available, we upgraded the thermometer and thermocouples. While that upgrade enhanced precision, it did not necessarily improve the overall function of the TIC. There is a wide range of thermometers available that can fit almost any research budget. Another advanateg of our modlar design is that should a researcher lack the expertise or tools to test and solder component connections as we did, all of these components are available in versions that could be run from battery power rather than hardwired into the GC.

One of the criticisms of *CT*_max_ has been the partially subjective nature of determining LOE (Lutterschmidt and Hutchison 1997; Chown and Nicolson 2004; Ørsted et al, 2018). We believe that the TIC improves our ability to determine when LOE has occurred in some insects. As the vials rotate, the insect must maintain enough coordination to keep themselves attached to the glass walls of the vial. Once they reach their *CT*_max_ they begin to fall and are typically unable to right themselves. The constant rotation may lead to the insect righting itself, but once the vial was upside down, they were unable to maintain their position and *CT*_max_ was recorded. We acknowledge this may introduce some error to the estimation of *CT*_max_, however, the magnitude of the error would be equivalent to the ramp rate of the oven and the RPM of the carousel. For example, at two RPM and 0.5 °C · min^-1^ the insect would fall from the walls twice per min if they were unable to maintain enough coordination to grip the walls. If the observer were to wait until the second fall this would lead to a recorded *CT*_max_ within 0.25 °C of the actual *CT*_max_ of the subject. We believe this still produces a reliable and repeatable estimate. Furthermore, we are currently investigating the incorporation of camera equipment that can be affixed to the TIC which can record activity throughout an experimental run. This could allow repeated scoring of LOE to further remove observational bias in the system.

We presented two proof-of-concept studies here as well. The first was a dynamic test of the *CT*_max_ of *C. vittatus*. We found that this beetle appears to have a high tolerance to heat. The field temperatures immediately preceding their collection were, on average, 30 °C cooler than their *CT*_max_ of 52.1 °C. This suggests that the insects are pre-acclimated for high heat tolerance, which makes sense given that during the cotton growing season in AZ, field temperatures can exceed 48 °C. We also found that acute exposure to high heat stress reduced the overall consumption of prey items after a short recovery period. Such a response has previously been observed in other predators such as *Serangium japonicum* (Yao et al, 2019), early stage *Aphidoletes aphidimyza* (Wang et al, 2022), among others (Bai et al, 2022; Tscholl et al, 2022; Martinez et al, 2023). Further testing would be needed to determine if additional time would be needed to fully recover from such extreme heat exposure.

In conclusion, while the cost for laboratory equipment can be particularly prohibitive for ecologists in resource poor situations, we believe the repurposing of free or low-cost laboratory equipment can lower the barrier to entry for studying thermal biology. While we have dubbed our system the Thermal Insect Carousel, any terrestrial animal that can be placed in a scintillation vial could be assayed in a comparable device. The modularity of the TIC further enhances its utility across ecological studies. We encourage researchers to explore similar innovations, fostering both accessibility and environmental stewardship in scientific infrastructure.

## Supporting information

TIC Stability Test data

Collops Stress Test data

## ACKNOWLEDGEMENTS

Mention of trade names or commercial products in this publication is solely to provide specific information and does not imply recommendation or endorsement by the United States Department of Agriculture. USDA is an equal opportunity provider and employer.

## AUTHOR CONTRIBUTION STATEMENT

**Gabriel Zilnik**: Conceptualization; Data Curation; Formal Analysis; Investigation; Methodology; Project Administration; Resources; Supervision; Validation; Visualization; Writing – Original Draft Preparation; Writing – Review & Editing. **Paul V. Merten**: Conceptualization; Investigation; Methodology; Resources; Supervision; Validation; Writing – Review & Editing. **Miles T. Casey**: Data Curation; Investigation; Methodology; Visualization; Writing – Review & Editing. **Scott A. Machtley**: Data Curation; Investigation; Methodology; Supervision; Writing – Review & Editing. **James R. Hagler**: Conceptualization; Investigation; Methodology; Project Administration; Resources; Supervision; Validation; Writing – Original Draft Preparation; Writing – Review & Editing.

## Data Availability Statement

All data presented in this paper is available in the supplementary materials and upon request from the corresponding author.

## Notes

### Competing Interest Statement

The authors have declared no competing interest.

